# Molecular and connectomic vulnerability shape cross-disorder cortical abnormalities

**DOI:** 10.1101/2022.01.21.476409

**Authors:** Justine Y. Hansen, Golia Shafiei, Jacob W. Vogel, Kelly Smart, Carrie E. Bearden, Martine Hoogman, Barbara Franke, Daan van Rooij, Jan Buitelaar, Carrie R. McDonald, Sanjay M. Sisodiya, Lianne Schmaal, Dick J. Veltman, Odile A. van den Heuvel, Dan J. Stein, Theo G. M. van Erp, Christopher R. K. Ching, Ole A. Andreassen, Tomas Hajek, Nils Opel, Gemma Modinos, André Aleman, Ysbrand van der Werf, Neda Jahanshad, Sophia I. Thomopoulos, Paul M. Thompson, Richard E. Carson, Alain Dagher, Bratislav Misic

## Abstract

Numerous brain disorders demonstrate structural brain abnormalities, which are thought to arise from molecular perturbations or connectome miswiring. The unique and shared contributions of these molecular and connectomic vulnerabilities to brain disorders remain unknown, and has yet to be studied in a single multi-disorder framework. Using MRI morphometry from the ENIGMA consortium, we construct maps of cortical abnormalities for thirteen neurodevelopmental, neurological, and psychiatric disorders from *N* = 21 000 patients and *N* = 26 000 controls, collected using a harmonized processing protocol. We systematically compare cortical maps to multiple micro-architectural measures, including gene expression, neurotransmitter density, metabolism, and myelination (molecular vulnerability), as well as global connectomic measures including number of connections, centrality, and connection diversity (connectomic vulnerability). We find that regional molecular vulnerability and macroscale brain network architecture interact to drive the spatial patterning of cortical abnormalities in multiple disorders. Local attributes, particularly neurotransmitter receptor profiles, constitute the best predictors of both disorder-specific cortical morphology and cross-disorder similarity. Finally, we find that cross-disorder abnormalities are consistently subtended by a small subset of network epicentres in bilateral sensory-motor, medial temporal lobe, precuneus, and superior parietal cortex. Collectively, our results highlight how local biological attributes and global connectivity jointly shape cross-disorder cortical abnormalities.

## INTRODUCTION

The brain is a network with intricate connection patterns among individual neurons, neuronal populations, and large-scale brain regions. The wiring of the network supports propagation of electrical signals, as well as molecules needed for growth and repair. This complex system is vulnerable to multiple neurological, psychiatric and neurodevelopmental disorders. Pathological perturbations—including altered cellular morphology, cell death, aberrant synaptic pruning and miswiring— disrupt inter-regional communication and manifest as overlapping groups of sensory, motor, cognitive and affective symptoms. How different disorders are shaped by local and global vulnerability is unknown.

Indeed, several studies have demonstrated cross-disorder connectomic vulnerability, where regions and white matter pathways are targeted non-randomly. In particular, regions that are highly connected and potentially important for communication tend to be disproportionately affected by disease [26, 114]. A similar phenomenon is observed for connections that support multiple communication pathways [28]. In neurodegenerative diseases such as Alzheimer’s and Parkinson’s diseases, emerging evidence suggests pathological misfolded proteins spread trans-synaptically, such that the connectivity of the brain shapes the course and expression of these diseases [56, 84, 90, 91, 101, 125, 128, 134]. Recent evidence also suggests that patterns of tissue volume loss in schizophrenia are circumscribed by structural and functional connection patterns [103, 124]. Collectively, these studies demonstrate that both neurodevelopmental and neurodegenerative brain diseases are influenced by network connectivity [40, 115].

The effects of disease can also be driven by local cellular and molecular vulnerability. Namely, local patterns of gene expression [4, 18, 75], neurotransmitter receptor profiles [53], cellular composition [100], and metabolism [15, 16, 122, 123] may predispose individual regions to stress and, ultimately, pathology. Importantly, local and global vulnerability are not necessarily mutually exclusive; some diseases may originate from local pathologies that spread selectively along the network to other vulnerable regions. How local attributes and global connectivity shape cross-disorder pathology remains an open question.

Here we map local molecular attributes (“molecular vulnerability”) and global network connectivity (“connectomic vulnerability”) to case versus control cortical thickness abnormalities of thirteen different neurological, psychiatric, and neurodevelopmental diseases and disorders from the ENIGMA consortium [111]. We consistently find that disorder-specific cortical abnormality is shaped more by the local molecular fingerprints of brain regions than network embedding. Interestingly, for disorders that are better predicted by molecular attributes, we find that the spatial patterning of cortical abnormalities reflects the underlying network architecture, suggesting that the local molecular and global connectomic contributions to disorder effects may interact.

Next, we study cross-disorder similarity and find that regions with similar molecular make-up tend to be similarly affected across disorders. Collectively, the present report highlights how local and global factors interact to shape cross-disorder cortical morphology.

## RESULTS

We collected thirteen spatial maps of cortical abnormalities from the ENIGMA consortium for the following diseases, disorders, and conditions: 22q11.2 deletion syndrome (22q) [109], attention-deficit/hyperactivity disorder (ADHD) [55], autism spectrum disorder (ASD) [119], idiopathic generalized epilepsy [127], right temporal lobe epilepsy [127], left temporal lobe epilepsy [127], depression [99], obsessive-compulsive disorder (OCD) [13], schizophrenia [117], bipolar disorder (BD) [51], obesity [82], schizotypy [62], and Parkinson’s disease (PD) [63]. For simplicity, we refer to diseases, disorders, and conditions as “disorders” throughout the text. While most disorders show decreases in cortical thickness, some (e.g. 22q, ASD, schizotypy) also show regional increases in cortical thickness. We therefore refer to the cortical measure as “cortical abnormalitiy”. All cortical abnormality maps were collected from adult patients, following identical processing protocols, for a total of over 21 000 scanned patients against almost 26 000 controls. To assess the extent to which each abnormality pattern is informed by molecular attributes and network connectivity, we defined a lar fingerprint of a region was defined using the gene expression gradient, neurotransmitter receptor gradient, excitatory-inhibitory receptor density ratio, glycolytic index, glucose metabolism, synapse density, and myelination (Fig. 1a). Likewise, we defined the connectivity fingerprint of a region by calculating the strength, betweenness centrality, closeness centrality, mean Euclidean distance, participation coefficient, clustering coefficient, and mean first passage time of a weighted structural connectivity matrix from 70 healthy adults (Fig. 1b; see *Methods* for details). Collectively, these graph measures aim to capture the connectedness, centrality, and connection diversity of regions in the network. All analyses were conducted using the 68-region Desikan Killiany parcellation [19, 29], as this is the native and only available representation of ENIGMA datasets.

**Figure 1.**
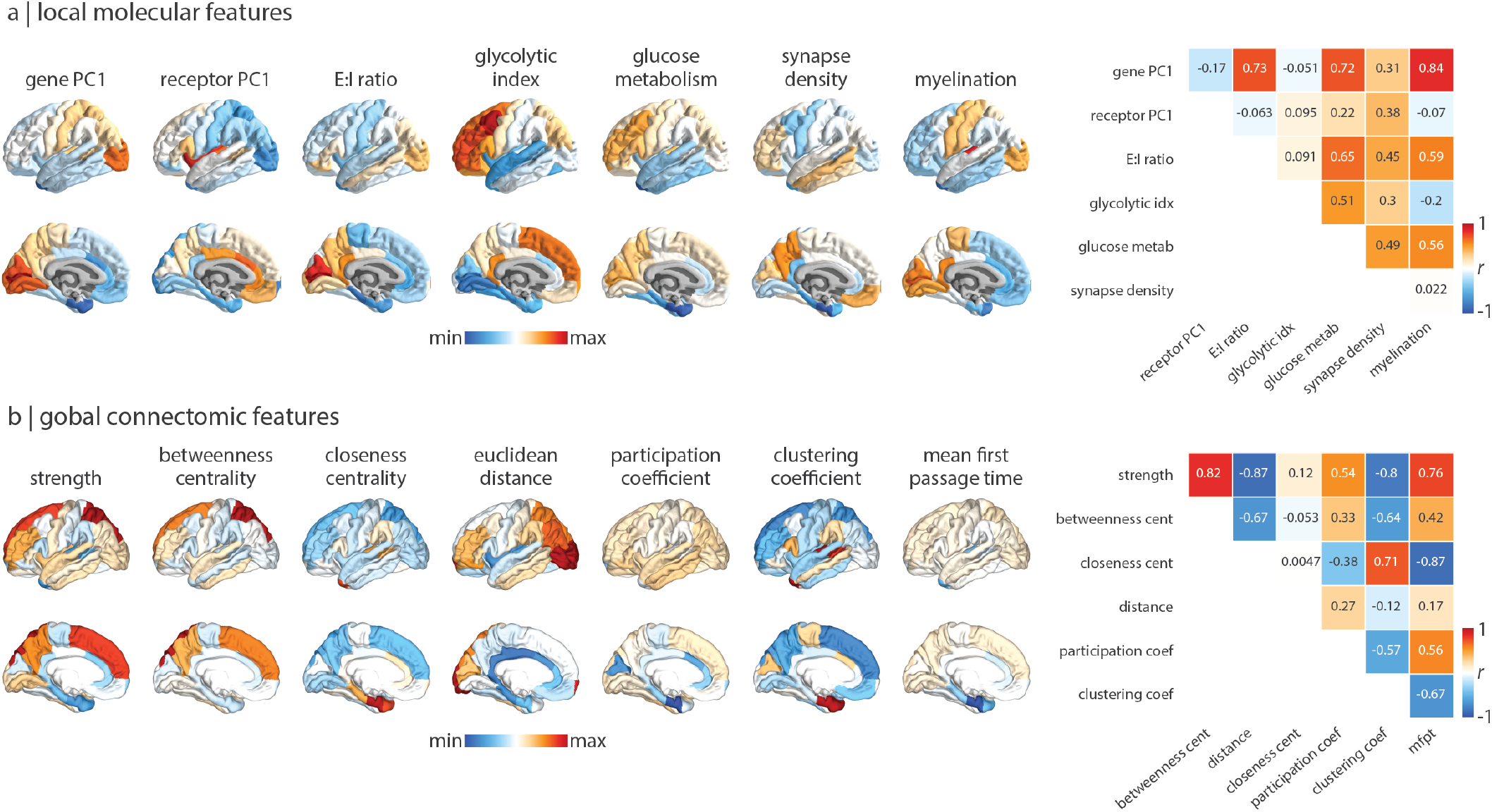
Molecular and connectomic cortical profiles. (a)–(b) Brain surfaces show the z-scored molecular (a) and connectomic (b) predictors used in the multilinear regression models. Heatmaps on the right show Pearson’s correlation coefficients between pairs of features. See *Methods* for details on how each feature was derived. Molecular predictors: gene PC1 = first component of 11 560 genes’ expression; receptor PC1 = first component of 18 PET-derived receptor/transporter density; E:I ratio = excitatory:inihibitory receptor density ratio; glycolytic index = amount of aerobic glycolysis; glucose metabolism = [^18^F]-labelled fluorodeoxyglucose (FDG) PET image; synapse density = synaptic vesicle glycoprotein 2A (SV2A)-binding [^11^*C*]UCB-J PET tracer; myelination = T1w/T2w ratio. Connectivity predictors: strength = sum of weighted connections; betweenness = fraction of all shortest paths traversing region *i*; closeness = mean shortest path length between region *i* and all other regions; Euclidean distance = mean Euclidean distance between region *i* and all other regions; participation coefficient = diversity of connections from region *i* to the seven Yeo-Krienen resting-state networks [130]; clustering = fraction of triangles including region *i*; mean first passage time = average time for a random walker to travel from region *i* to any other region.

### Local and global contributions to disorder-specific cortical morphology

To assess the extent to which cortical abnormalities of all thirteen disorders are informed by molecular gradients versus measures of network connectivity, we fit a multilinear model between molecular or connectivity predictors and abnormality maps for each disorder separately, for a total of 13 × 2 = 26 model fits (Fig. 2a; for results when measures of network connectivity were computed on the binary structural connectome and the functional connectome, see Fig. S1; for results when molecular and connectivity fingerprint at each brain region. The molecumolecular and connectomic predictors are combined, see Fig. S2). Next, we conducted a dominance analysis for each multilinear model [5, 17, 67, 102]. Dominance analysis distributes the 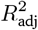 across input variables as a measure of contribution (“dominance”) that each input variable has on the cortical thinning pattern (Fig. 2b). Dominance was assessed against a spatial autocorrelation-preserving null model (“spin test”, see *Methods* for details), and each model was cross-validated in a distance-dependent manner (Fig. S3; [46]).

**Figure 2.**
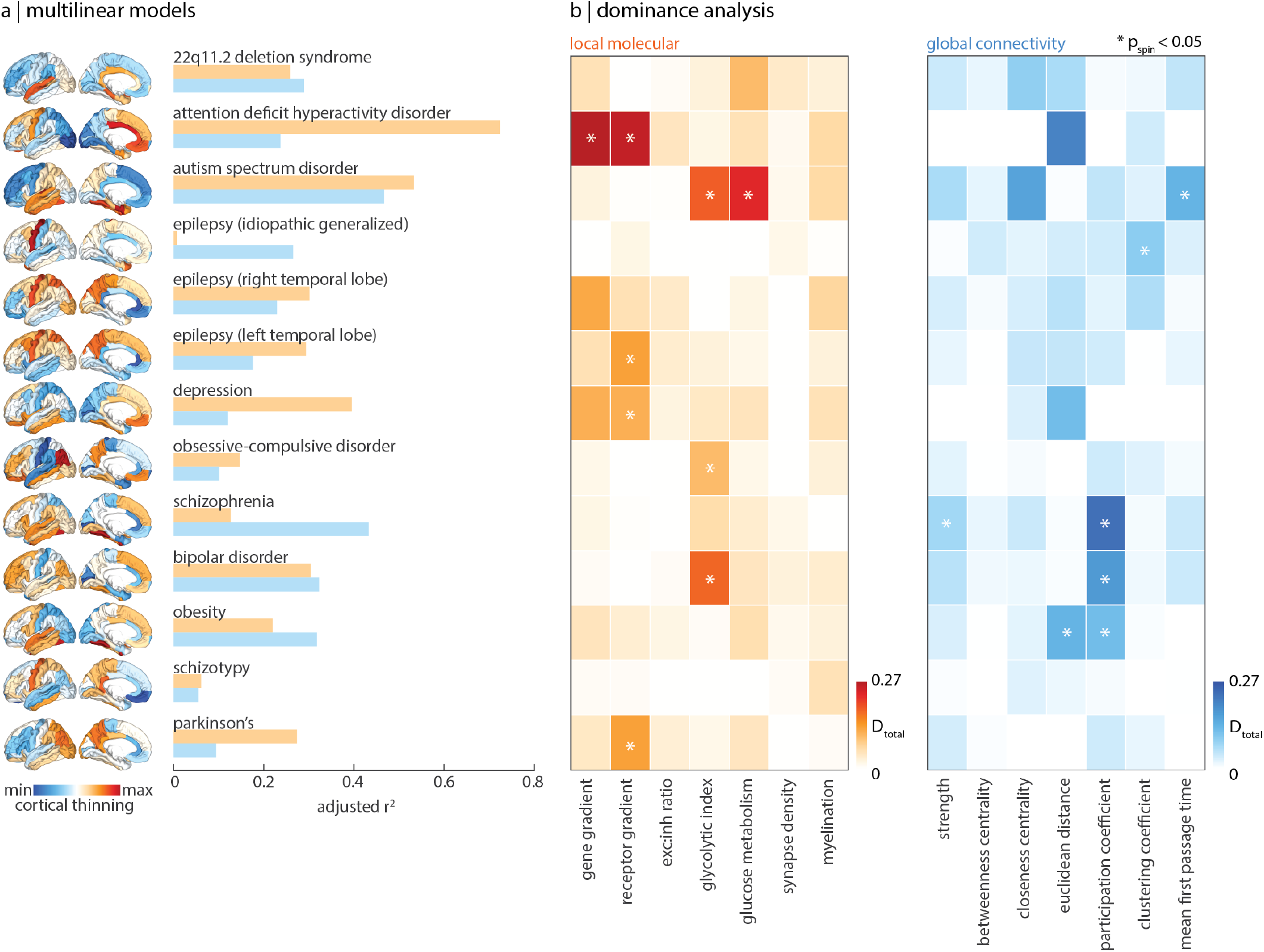
Local and global contributions to disorder-specific cortical morphology. (a) A total of twenty-six multilinear models were fit between local molecular and global connectome predictors to cortical abnormality maps of thirteen different disorders (surface plots, left). Adjusted *R*^2^ is shown in the bar plot (orange: molecular; blue: connectivity). (b) Dominance analysis was applied to assess the contribution of each input variable (done separately for molecular (orange) and connectivity (blue) predictors) to the fit of the model. Significance was assessed using the spin-test (two-tailed); asterisks represent *p*_spin_ < 0.05.

We find that the fit between molecular predictors and cortical thinning is greater than that between connectivity predictors and cortical thinning for most disorders (Fig. 3). Notably, the variance in cortical thickness of schizotypy (a possible precursor of schizophrenia that is poorly defined in the brain [66]) and idiopathic generalized epilepsy (a form of epilepsy that is thought to be informed by genetics instead of brain structural abnormalities [24, 35]) are poorly explained by both biological gradients and network measures of the brain. On the other hand, ADHD, ASD, OCD, PD, and depression are better predicted by biological predictors, whereas schizophrenia, 22q deletion syndrome, and bipolar disorder are better predicted by connectivity predictors (Fig. 3).

**Figure 3.**
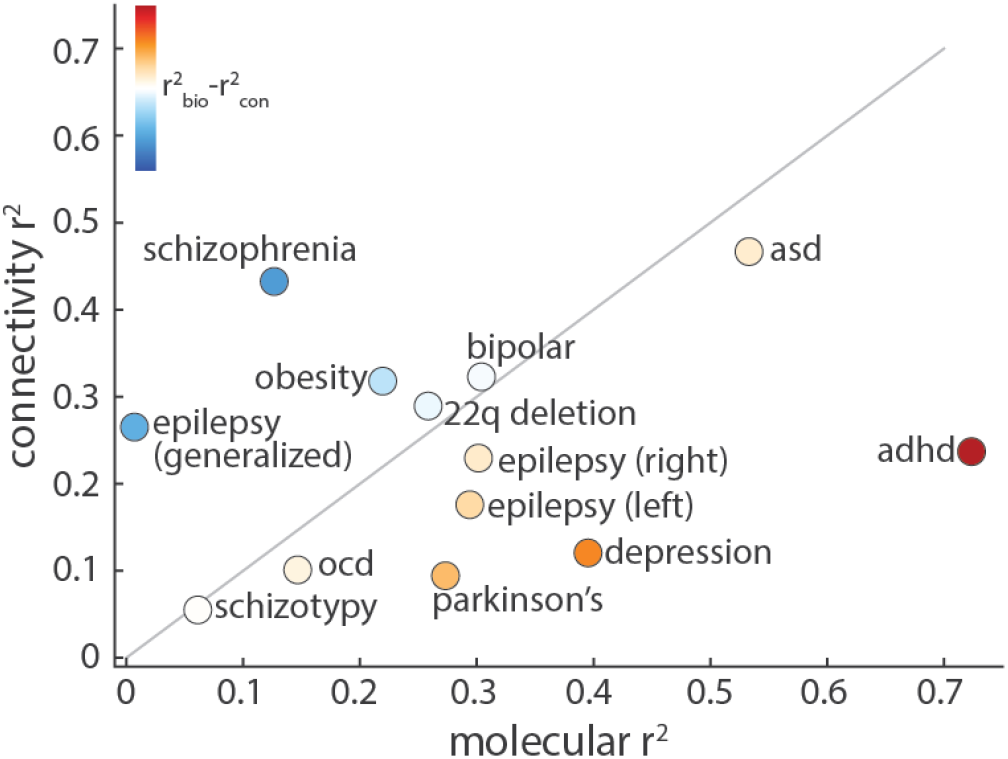
Comparing molecular and connectomic contributions to disorder-specific cortical differences. The local molecular 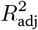 of each disorder is plotted against the global connectivity 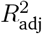. The grey line indicates the identity line and circle colour represents the difference between molecular and connectomic 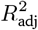, such that warm colours represent disorders that are better predicted by molecular predictors, and cool colours represent disorders that are better predicted by connectomic predictors.

From the dominance analysis, we find that certain predictors are consistently unimportant. Indeed, synapse density and myelination demonstrate less dominance than microscale gradients such as the gene expression gradient, neurotransmitter receptor gradient, and metabolic gradients. Connectivity predictors, particularly measures of centrality, demonstrate less dominance than more fundamental measures of connectivity such as number of connections (strength), distance, and connection diversity (participation coefficient). For completeness, we tested a third family of predictors related to temporal dynamics: magnetoencephalography (MEG)-derived power spectral densities for six canonical frequency bands (Fig. S4). However, no temporal predictors were significantly dominant toward any disorders so we excluded the temporal predictors from further analyses.

### Interactions between local and global vulnerability

The previous section separately addresses molecular and connectomic contributions to disease-specific cortical abnormalities. However, molecular attributes likely interact with network connectivity to shape disease pathology. These molecular mechanisms include gene expression, neurotransmitter expression, and metabolic pathways in the cell. In neurodegenerative diseases, this interaction may result in synaptic pruning and cortical atrophy whereas in neurodevelopmental disorders, the pathology may manifest as perturbations in network wiring during the embryonic stage [30]. We hypothesized that abnormalities in such molecular mechanisms at the regional level may spread trans-synaptically between connected regions, resulting in connectome-informed changes in cortical morphology that reflect an interplay between local vulnerability and network structure. For instance, two regions may both participate in many connections (have high degree), but one may be connected to more regions with local vulnerability. Thus, despite the fact that their connectomic profiles are similar, one region may have greater disease exposure than the other [14, 134].

To test the hypothesis that a region’s cortical thickness is driven by “exposure” to abnormalities of connected regions, we measured the extent to which disorders demonstrate network spreading patterns of cortical morphology [23, 101, 103]. The extent to which a disorder displays network-informed cortical changes is defined as the correlation between regional abnormality and mean abnormality of structurally connected neighbours (Fig. 4a). Importantly, significance was assessed using a spatial autocorrelation preserving null model to control for the effect of distance on cortical abnormality patterns. We also test the hypothesis that this network-spreading effect is functionally informed, whereby the cortical thickness of structurally connected neighbours is weighted by the functional connectivity between regions when calculating the mean (Fig. 4b; see *Methods* for details and Fig. S5 and S6 for scatter plots of regional abnormality versus mean neighbour abnormality across all thirteen disorders). We find that multiple disorders display a significant correlation between regional thickness and thickness of connected neighbours (0.23 *< r <* 0.80), suggesting that spatial patterning of disorders reflects the connection patterns between brain regions, above and beyond the effect of spatial autocorrelation (Fig. S5).

**Figure 4.**
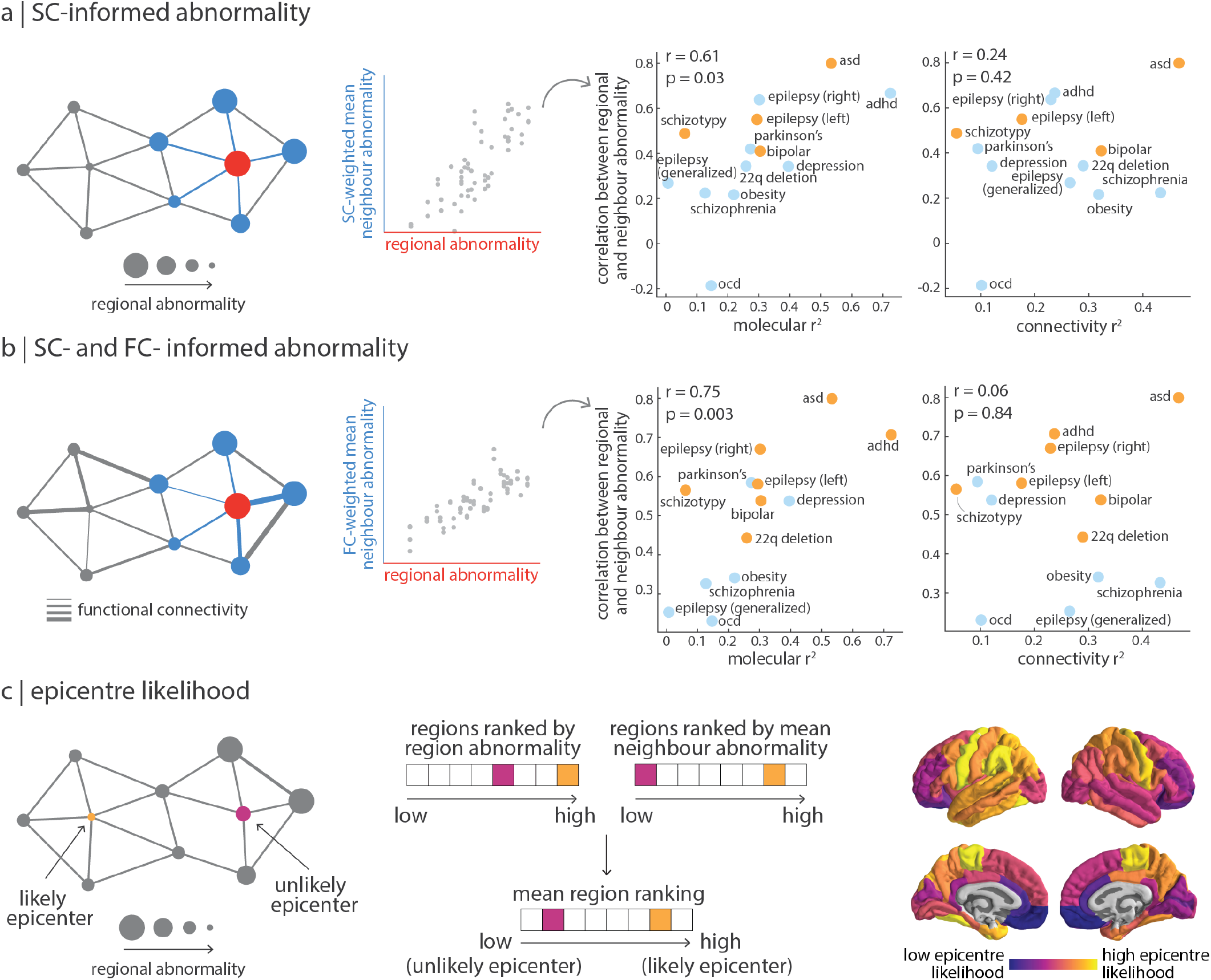
Interactions between molecular and connectomic vulnerability. (a) Left: schematic of structural connectivity informing disorder-related cortical changes. The correlation between SC-weighted mean neighbour abnormality and region abnormality represents the extent to which a disorder demonstrates network spreading disorder-specific cortical morphology. Right: this correlation coefficient was then correlated to both local molecular (left) and global connectivity (right)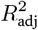. Yellow points refer to disorders where the correlation between region abnormality and SC-weighted mean neighbour abnormality is significant (*p*_spin_ < 0.05) (b) Left: likewise, mean neighbour abnormality can be additionally weighted by functional connectivity between regions. Right: correlation between the extent to which a disorder demonstrates SC- and FC-informed network spreading cortical morphology and local molecular (left) and global connectivity (right) 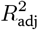. Yellow points refer to disorders where the correlation between region abnormality and SC- and FC-weighted mean neighbour abnormality is significant (*p*_spin_ < 0.05) (c) Left: a region with high abnormality that is also connected to regions with high abnormality is considered a likely disorder epicentre. Middle: epicentre likelihood was calculated as the mean rank of region and neighbour abnormality. Right: mean epicentre likelihood was calculated for all seven disorders that show a significant correlation between regional and neighbour abnormality.

Does molecular or connectomic predictability of a disorder pattern (Fig. 2a) relate to network spreading? Interestingly, the extent to which a disorder can be predicted from molecular attributes (i.e. yellow *R*^2^ in Fig. 2a) is positively correlated with the extent to which a disorder displays evidence of network spreading (*r* = 0.61, *p* = 0.03 when weighted by SC only as shown in Fig. 4a; *r* = 0.75, *p* = 0.003 when weighted by FC and SC as shown in Fig. 4b). Notably, we do not observe this relationship with the extent to which a disorder can be predicted from global connectivity (i.e. blue *R*^2^ in Fig. 2a; *r* = 0.24, *p* = 0.42 when weighted by SC only, Fig. 4a; *r* = 0.06, *p* = 0.84 when weighted by FC and SC, Fig. 4b). In other words, for disorders with cortical morphologies that more strongly depend on molecular attributes, we also observe a greater effect of disorder exposure. Although we previously found that the cortical patterning of a disorder is less influenced by network embedding per se (e.g. centrality or connection diversity), here we show that it is instead more influenced by network-driven exposure to regions with local vulnerability. This finding is significant because it shows that connectome architecture interacts with local vulnerability.

Brain regions with high abnormality and high neighbour abnormality are likely to act as an epicentre of the network spreading disorder pattern, since the region is both heavily affected and facilitates the spread of atypical morphology [14, 103, 133]. We calculated epicentre likelihood of each brain region as the mean rank of regional and neighbour abnormality, such that regions with high node and neighbour abnormality would be labeled as likely epicentres (Fig. 4c). The measure identifies “disorder hubs”—regions that are both vulnerable to disorder-specific changes but also embedded in a highly atypical network cluster. Epicentre likelihood was only calculated for brain maps with significant correlation between their node and neighbour abnormality (network spreading disorders). This list comprised of: 22q11.1 deletion syndrome, ADHD, ASD, right and left temporal epilepsy, bipolar disorder, and schizotypy (Fig. S7). We averaged epicentre likelihood of these seven disorders to produce a map of epicentre likelihood across disorders that demonstrate network spreading disorder-specific cortical morphology (Fig. 4c, right). We find that cross-disorder epicentre likelihood is highest in bilateral sensory-motor cortex, medial temporal lobe, precuneus, and superior parietal cortex.

### Brain regions with similar molecular annotations are similarly affected across disorders

In the previous sections, we mapped molecular annotations and network measures to each disorder separately. Here, we focused on disorder similarity. For every region we constructed a 13-element vector of abnormality values, where each element corresponds to cortical change in that region in one disorder. We then correlated regional vectors with each other to estimate how similarly two regions are affected across the thirteen disorders (Fig. 5a).

**Figure 5.**
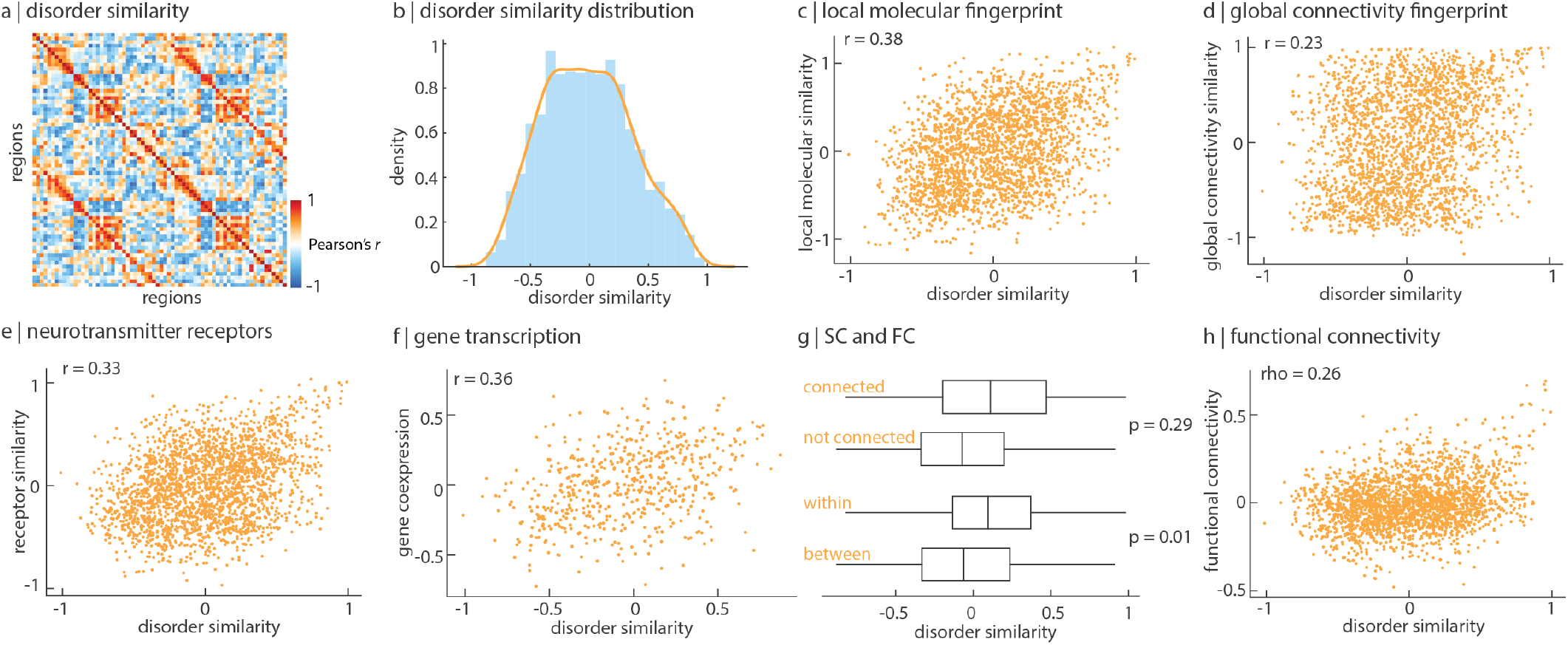
Brain regions with similar molecular annotations are similarly affected across disorders. (a) Disorder similarity was computed as the pairwise correlation of regional cortical thickness across all thirteen disorders such that pairs of regions with high disorder similarity are similarly affected across disorders. (b) The upper triangle of the disorder similarity matrix is approximately normally distributed. (c) Disorder similarity is significantly correlated to molecular attribute similarity, after distance-regression (*r* = 0.38, *p <* 0.001). (d) Disorder similarity is less correlated to global connectome attribute similarity after distance-regression (*r* = 0.22, *p <* 0.001). (e) Disorder similarity is significantly correlated to neurotransmitter receptor similarity, after distance-regression (*r* = 0.33, *p <* 0.001). (f) Left hemisphere disorder similarity is significantly correlated to gene coexpression, after distance-regression (*r* = 0.36, *p <* 0.001). (g) Disorder similarity is significantly greater within intrinsic functional networks than between networks, against the spin test (*p* = 0.005; top). Disorder similarity is significantly greater between structurally connected regions than regions that are not connected, against a degree- and edge-length-preserving null model (*p* = 0.003 [11]). (h) Disorder similarity is significantly correlated to functional connectivity, after distance-regression (*r* = 0.26, *p <* 0.001).

We first asked whether brain regions with similar molecular versus connectivity fingerprints show greater disorder similarity. Molecular similarity was likewise computed as the pairwise regional correlation of molecular predictors, and vice versa for connectivity. To account for spatial autocorrelation in molecular and connectomic attributes, we regressed the exponential trend with Euclidean distance out of molecular similarity, connectivity similarity, and disorder similarity (Fig. S8a, b, c). After distance-regression, we find that disorder similarity is significantly correlated with molecular similarity (*r* = 0.38, *p <* 0.001; Fig. 5c). On the other hand, the correlation between distance-regressed connectivity similarity and disorder similarity was smaller and less visually apparent, albeit significant (*r* = 0.23, *p <* 0.001; Fig. 5d).

Two of the molecular predictors included in the present report are summary measures of much more expansive molecular annotations: the gene expression gradient and the neurotransmitter receptor gradient. We therefore asked whether inter-regional similarity of these molecular attributes confers similar predisposition to disease. We computed gene coexpression and neurotransmitter receptor similarity matrices, regressed out the exponential trend with Euclidean distance as before (Fig. S8d, e, [39, 47]), and correlated these matrices with disorder similarity (Fig. 5e, f). We find a significant correlation between disorder similarity and neurotransmitter receptor similarity (*r* = 0.33, *p <* 0.001) as well as gene coexpression (*r* = 0.36, *p <* 0.001) [3, 47]. Altogether, our results indicate that regions with similar biological composition are similarly affected across disorders.

We finally ask whether disorder similarity might analogously be informed by structural and functional connectivity between regions. We compared the disorder similarity matrix to weighted structural and functional connectomes. First, we find that brain regions that are structurally connected are more likely to change similarly across disorders than regions that are not structurally connected, although this result is non-significant against a degree and edge-length preserving null model (Fig. 5g [11]; see *Null models*). Second, we find that brain regions that are within the same intrinsic functional network are more likely to change similarly than regions between functional networks, against the spin-test (Fig. 5g, *p*_spin_ = 0.01). Finally, we find a positive significant correlation between disorder similarity and functional connectivity (*r* = 0.26, *p*_spin_ = 0.002; Fig. 5h). Consistent with the previous subsection, these results collectively suggest that areas that share molecular attributes and connections are similarly affected across disorders.

## DISCUSSION

In the present report, we comprehensively map local molecular attributes and global measures of connectivity to the cortical morphology of thirteen different neurological, psychiatric, and neurodevelopmental disorders. We consistently find that local attributes govern both disorder-specific abnormalities and cross-disorder similarity more than global connectivity and regional dynamics. In addition, we find that molecular mechanisms interact with the structural and functional architectures of the brain to guide cross-disorder abnormality patterns. Altogether, our results highlight how molecular and connectomic vulnerability shape cross-disorder cortical abnormalities.

This work builds on a growing literature on cross-disorder effects, and how shared vulnerability may potentially transcend traditional diagnostic boundaries [28, 61, 120, 129]. It is becoming increasingly clear that pathology is governed by layers of abnormal processes, at the molecular and cellular level, to neural dynamics, to large-scale brain networks. Aligning high-quality maps of disorder-specific cortical changes to a common reference frame of local and global attributes allows us to systematically relate the effect of disease to multiple scales of organization. By taking a cross-modal and cross-disorder approach we reveal that, despite different clinical presentation and label, there exists some commonality across diseases including predictors that are ubiquitously important as well as interplay between local vulnerability and network structure.

Interestingly, we find that the principal gradient of receptor distribution is particularly dominant towards disease-specific cortical morphology. This receptor gradient represents the maximal variance of density distributions from fourteen receptors and four transporters across nine different neurotransmitter systems, and therefore captures how brain regions may integrate exogenous signals differently [47, 104]. This gradient is a powerful predictor of ADHD, and is the only significantly dominant predictor of left temporal lobe epilepsy, depression, and Parkinson’s disease. Indeed, neurotransmitter dysfunction is thought to underlie multiple disorders, including dopamine release in PD and schizophrenia or serotonin reuptake in depression. Modern therapeutics are designed to selectively manipulate neurotransmitter function for the purpose of alleviating behavioural symptoms, as opposed to altering brain structure. Our findings confirm the fundamental contribution of neurotransmitters to a wide spectrum of diseases, but they also highlight an important link between the spatial patterning of neurotransmitter receptors and cortical disorder morphology itself [47].

We generally find that cortical abnormality is better predicted by local vulnerability compared to global connectomic vulnerability. One possible reason for the relatively poorer performance of connectivity predictors is that they are generic measures of a region’s embedding in a network (number of connections, centrality, connection diversion) but do not consider how this embedding exposes regions to pathology elsewhere in the network. Indeed, we find that disorders whose cortical morphology is better reflected by local vulnerability also bear a prominent signature of network architecture (e.g. ASD, ADHD, 22q, temporal lobe epilepsy, schizotypy, bipolar disorder). Namely, in these disorders, areas with greater change are disproportionately more likely to be structurally- and functionally-connected with each other. This suggests a network spreading phenomenon where focal pathology or perturbation propagates to connected regions, resulting in cortical abnormality that is correlated with the underlying connection patterns [40]. This interaction between local vulnerability and connectomic vulnerability has previously been reported in neurodegenerative syndromes where the trans-synaptic spreading of misfolded proteins appears to be guided and amplified by local gene expression [25, 50, 92].

The interaction between molecular vulnerability and network structure naturally raises the question of what are the network epicentres of cortical disorder maps. We find epicentres—regions with high abnormality that are also strongly connected with other regions with high abnormality—in the sensory-motor cortex, medial temporal lobe, precuneus, and superior parietal cortex. That the sensory-motor cortex is an epicentre is consistent with recent reports that multiple psychiatric and neurological disorders are accompanied by sensory deficits and reduced motor control [10, 57, 69]. Indeed, the sensorymotor cortex has been previously established as a functional hub in temporal lobe epilepsies and across multiple psychiatric disorders [61, 65]. Interestingly, both the bilateral precuneus and superior parietal cortex are members of the brain’s putative rich club—densely interconnected regions that are thought to support the integration and broadcasting of signals [113]. Rich club regions undergo changes in connectivity patterns in multiple diseases such as schizophrenia, Alzheimer’s, and Huntington’s [28, 115, 116]. We complement this work by showing that the precuneus and superior parietal cortex are both vulnerable to cortical abnormality and, by virtue of their network embedding, increase disease exposure to connected regions. Conversely, although the anterior cingulate cortex (ACC) is implicated across multiple psychiatric disorders [45, 103], we do not find that the ACC is an epicentre of cross-disorder cortical morphology. This suggests that although the ACC demonstrates considerable local vulnerability in a subset of brain disorders, it is not consistently involved across the seven disorders included in the epicentre analyses. Altogether, despite variable cortical morphology patterns across the thirteen disorders, when looked at through the lens of network connectivity, we see a more consistent and compact subset of potential epicentres, suggesting greater commonality among diseases than previously appreciated.

The present work should be considered along some important methodological considerations. First, although the ENIGMA consortium standardizes preprocessing pipelines and provides large *N* datasets, allowing for robust results and meaningful comparison between disorder-specific cortical abnormality maps, working with ENIGMA data also has caveats: (1) the measures of cortical abnormality are effect sizes between patients and controls and do not represent tissue volume loss/gain, (2) some of the patient populations included have co-morbidities and patients may be undergoing treatment, and (3) all analyses were conducted at the level of 68 cortical brain areas, limiting regional specificity and precluding analyses of the subcortex and cerebellum. Second, despite the fact that structural connectomes were reconstructed from high resolution diffusion spectrum imaging, diffusion tractography is still prone to false-positives and false-negatives [58, 68, 132]. Third, both local biological and global connectivity predictors are derived from state-of-the-art open-access datasets in healthy participants, but they do not capture individual variability or changes across the lifespan—both of which are key factors in neurological, psychiatric, and neurodevelopmental disorders. Additionally, the biological predictors are limited by imaging modality and, in the case of the gene and receptor gradients, by the subset of genes and receptors included in the data decomposition. Fourth, we assessed contribution of multiple predictors to disorder maps using simple but robust linear models that are not sensitive to non-linear contributions or higher-order interactions among the predictors. Fifth, the linear models used in the present analyses assume independence between observations, which is not the case in the brain; we therefore employ spatial-autocorrelation preserving null models to account for the spatial dependencies between regions throughout the report. Finally, although the present report spans a wide range of neurological, psychiatric, and neurodevelopmental disorders, results are only valid for this subset of disorders. Future work is needed to map local and global vulnerabilities to the many more brain diseases and disorders that exist.

In summary, we find that molecular and connectomic vulnerability jointly shape cross-disorder cortical abnormalities. Cross-disorder regional vulnerability is largely driven by molecular fingerprints, including neurotransmitter receptor densities and gene expression, while connection patterns among vulnerable regions further amplify this vulnerability. Our results highlight how an integrative, multi-modal approach can illuminate the contributions of local biology and connectome architecture to brain disease.

## METHODS

All code and data used to perform the analyses can be found at https://github.com/netneurolab/hansen_crossdisorder_vulnerability. Molecular predictors can also be found in the neuromaps toolbox (https://netneurolab.github.io/neuromaps/[71]).

### Cortical disorder maps

Patterns of cortical thickness were collected for the available thirteen neurological, neurodevelopmental, and psychiatric disorders from the ENIGMA consortium and the *enigma* toolbox (https://github.com/MICA-MNI/ENIGMA; [64]) including: 22q11.2 deletion syndrome (22q) [109], attention-deficit/hyperactivity disorder (ADHD) [55], autism spectrum disorder (ASD) [119], idiopathic generalized epilepsy [127], right temporal lobe epilepsy [127], left temporal lobe epilepsy [127], depression [99], obsessive-compulsive disorder (OCD) [13], schizophrenia [117], bipolar disorder (BD) [51], obesity [82], schizotypy [62], and Parkinson’s disease (PD) [63]. The ENIGMA (Enhancing Neuroimaging Genetics through Meta-Analysis) Consortium is a data-sharing initiative that relies on standardized image acquisition and processing pipelines, such that disorder maps are comparable [111]. Altogether, over 21 000 patients were scanned across the thirteen disorders, against almost 26 000 controls. The values for each map are z-scored effect sizes (Cohen’s *d*) of cortical thickness in patient populations versus healthy controls. Imaging and processing protocols can be found at http://enigma.ini.usc.edu/protocols/.

### Structural and functional data acquisition

Structural and functional data were collected at the Department of Radiology, University Hospital Center and University of Lausanne, on *n* = 70 healthy young adults (16 females, 25.3 ± 4.9 years). Informed consent was obtained from all participants and the protocol was approved by the Ethics Committee of Clinical Research of the Faculty of Biology and Medicine, University of Lausanne. The scans were performed in a 3-T MRI scanner (Trio; Siemens Medical), using a 32-channel head coil. The protocol included (1) a magnetization-prepared rapid acquisition gradient echo (MPRAGE) sequence sensitive to white/grey matter contrast (1 mm in-plane resplution, 1.2 mm slice thickness), (2) a DSI sequence (128 diffusion-weighted volumes and a single b0 volume, maximum b-value 8 000s/mm^2^, 2.2 × 2.2 × 3.0 mm voxel size), and (3) a gradient echo-planar imaging (EPI) sequence sensitive to blood-oxygen-level-dependent (BOLD) contrast (3.3 mm in-plane resolution and slice thickness with a 0.3 mm gap, TR 1 920 ms, resulting in 280 images per participant). Participants were not subject to any overt task demands during the fMRI scan. The Lausanne dataset is available at https://zenodo.org/record/2872624#.XOJqE99fhmM.

### Structural network reconstruction

Grey matter was parcellated according to the 68-region Desikan-Killiany cortical atlas [29]. Structural connectivity was estimated for individual participants using deterministic streamline tractography. The procedure was implemented in the Connectome Mapping Toolkit [27], initiating 32 streamline propagations per diffusion direction for each which matter voxel. Collating each individual’s structural connectome was done using a group-consensus approach that seeks to preserve the density and edge-length distributions of the individual connectomes [11].

We first collated the extant edges in the individual participant matrices and binned them according to length. The number of bins was determined heuristically, as the square root of the mean binary density across participants. The most frequently occurring edges were then selected for each bin. If the mean number of edges across participants in a particular bin is equal to *k*, we selected the *k* edges of that length that occur most frequently across participants. To ensure that interhemispheric edges are not underrepresented, we carried out this procedure separately for inter- and intrahemispheric edges. The binary density for the final whole-brain structural connectome was 24.6%. For the weighted structural connectome, edges were weighted by the log-transform of the mean non-zero streamline density, scaled to values between 0 and 1.

### Functional network reconstruction

Functional MRI data were preprocessed using procedures designed to facilitate subsequent network exploration [86]. fMRI volumes were corrected for physiological variables, including regression of white matter, cerebrospinal fluid, and motion (3 translations and 3 rotations, estimated by rigid body coregistration). BOLD time series were then subjected to a low-pass filter (temporal Gaussian filter with full width at half maximum equal to 1.92 s). The first four time points were excluded from subsequent analysis to allow the time series to stabilize. Motion “scrubbing” was performed as described by [86]. The data were parcellated according to the same 68-region Desikan-Killiany atlas used for the structural network. Individual functional connectivity matrices were defined as zero-lag Pearson correlation among the fMRI BOLD time series. A group-consensus functional connectivity matrix was estimated as the mean connectivity of pairwise connections across individuals. Note that one individual did not undergo an fMRI scan and therefore the functional connectome was composed of *n* = 69 participants.

### Biological predictors

A total of seven local biological predictors were used in the multilinear model to represent the influence that local biological attributes have on disorder-specific cortical morphology.

#### Gene expression gradient

The first principal component of gene expression (“gene gradient”) was used to represent the variation in gene expression levels across the left cortex. Gene expression data was collected by the Allen Human Brain Atlas as described in Hawrylycz et al. [49] and processed by *abagen*, an open-source Python toolbox [70]. A total of 11 560 genes with differential stability greater than 0.1 were retained in the region by gene matrix [48]. The left gene gradient was mirrored in the right hemisphere. A detailed account of the specific processing choices made can be found in Hansen et al. [46].

#### Receptor gradient

The first principal component of receptor density (“receptor gradient”) was used to represent the variation in receptor densities across the cortex. Receptor densities were estimated using PET tracer studies for a total of 18 receptors and transporters, across 9 neurotransmitter systems. These include dopamine (D1 [59], D2 [96, 105, 108, 131], DAT [33]), norepinephrine (NET [8, 22, 31, 95]), serotonin (5-HT1A [98], 5-HT1B [6, 41, 73, 76, 77, 85, 97, 98], 5-HT2A [9], 5-HT4 [9], 5-HT6 [87, 88], 5-HTT [9]), acetylcholine (*α*4*β*2 [6, 52], M1 [78], VAChT [1, 7]), glutamate (mGluR5 [32, 106]), GABA (GABAA [80]), histamine (H3 [42]), cannabinoid (CB1 [34, 79, 81, 93]), and opioid (MOR [60]). Volumetric PET images were registered to the MNI-ICBM 152 nonlinear 2009 (version c, asymmetric) template, averaged across participants within each study, then parcellated to 68 cortical regions. Parcellated PET maps were then z-scored before compiling all receptors/transporters into a region × receptor matrix of relative densities. Data were originally presented as an atlas in Hansen et al. [47].

#### Excitatory-inhibitory ratio

The excitatory-inhibitory ratio was computed as the ratio of z-scored PET-derived excitatory to inihibitory neurotransmitter receptor densities in the cortex, using the same dataset that was used to compute the receptor gradient. Excitatory neurotransmitter receptors included are: 5-HT2_A_, 5-HT_4_, 5-HT_6_, D_1_, mGluR_5_, *α*_4_*β*_2_, and M_1_. Inhibitory neurotransmitter receptors included are: 5-HT_1A_, 5-HT_1B_, CB_1_, D_2_, GABA_A_, H_3_, and MOR.

#### Glycolytic index

Aerobic glycolysis is the process of converting glucose to lactate in the presence of oxygen. It is traditionally calculated as the ratio of oxygen metabolism to glucose metabolism. Here, we use glycolytic index, a measure of aerobic glycolysis that mitigates certain limitations of using the traditional ratio [112]. Glycolytic index is defined as the residual after fitting glucose metabolism to oxygen metabolism in a linear regression model. Larger values indicate more aerobic glycolysis. Note that glycolytic index and the traditional ratio are highly correlated (see Vaishnavi et al. [112]). Data were collected, calculated, and made available by Vaishnavi et al. [112].

#### Glucose metabolism

Glucose metabolism in the cortex was measured in 33 healthy adults by administering [^18^*F*]-labelled fluorodeoxyglucose (FDG) for a PET scan, as described in detail in Vaishnavi et al. [112].

#### Synapse density

Synapse density in the cortex was measured in 76 healthy adults (45 males, 48.9 *±* 18.4 years of age) by administering [^11^*C*]UCB-J, a PET tracer that binds to the synaptic vesicle glycoprotein 2A (SV2A) [12, 20, 21, 36, 38, 54, 74, 83, 89, 107, 126]. Data were collected on an HRRT PET camera for 90 minutes post injection. Non-displaceable binding potential (BPND) was modelled using SRTM2, with the centrum semiovale as reference and 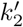 fixed to 0.027 (population value).

#### Myelination

Data from the Human Connectome Project (HCP, S1200 release) [43, 118] was used for measures of T1w/T2w ratios—a proxy for intracortical myelin—for 417 unrelated participants (age range 22– 37 years, 193 males), as approved by the WU-Minn HCP Consortium. Images were acquired on a Siemens Skyra 3T scanner, and included a T1-weighted MPRAGE sequence at an isotropic resolution of 0.7mm, and a T2-weighted SPACE also at an isotropic resolution of 0.7mm. Details on imaging protocols and procedures are available at http://protocols.humanconnectome.org/HCP/3T/imaging-protocols.html. Image processing includes correcting for gradient distortion caused by non-linearities, correcting for bias field distortions, and registering the images to a standard reference space. T1w/T2w ratios for each participant was made available in the surface-based CIFTI file format and parcellated into 68 cortical regions according to the Lausanne anatomical atlas [19].

### Connectivity predictors

A total of nine global connectome predictors were used in the multilinear model to represent the influence that global connectivity has on disorder-specific cortical morphology. In the main text, connectome measures were computed on the weighted structural connectome. Analyses were repeated using a binary structural connectome and an absolute functional connectome (Fig. S1). All connectivity measures were computed using the Python-equivalent of the Brain Connectivity Toolbox, *bctpy*.

#### Strength

The strength of region *i* is the sum of the edges connected to region *i*. For a binary structural connectome, the strength is equivalent to the degree, which is the number of links connected to region *i*.

#### Betweenness centrality

Betweenness centrality of region *i* is the fraction of all shortest paths between any two regions that traverse region *i*.

#### Closeness centrality

Closeness centrality is equivalent to the mean shortest path distance from region *i* to every other region in the network.

#### Euclidean distance

Mean Euclidean distance of a region to all other regions in the network represents how spatially close one region is to all other regions.

#### Participation coefficient

Participation coefficient was computed using the putative intrinsic functional networks of the brain [130]. Participation coefficient represents the connection diversity of a region. A region with high participation coefficient is well connected to several different networks, whereas a region with low participation coefficient primarily makes local (within-network) connections.

#### Clustering coefficient

The clustering coefficient of region *i* is the fraction of all triangles that are around region *i*. Equivalently, it is the fraction of all of region *i*’s neighbours that are also neighbours with each other. In the case of the weighted structural connectome, clustering coefficient is the average geometric mean of all triangles associated with the region.

#### Mean first passage time

The mean first passage time from region *i* to *j* is the expected amount of time it takes a random walker to reach region *j* from *i* for the first time. For each region, mean first passage time was averaged across regions, resulting in a mean mean first passage time representing the average amount of time it takes a random walker to travel from region *i* to any other region in the network for the first time.

### Temporal predictors

6-minute resting-state eyes-open magenetoencephalography (MEG) time-series were acquired from the Human Connectome Project (HCP, S1200 release) for 33 unrelated subjects (age range 22—35, 17 males) [43, 118]. Complete MEG acquisition protocols can be found in the HCP S1200 Release Manual. For each subject, we computed the power of the run at the vertex level across six different frequency bands: delta (2–4 Hz), theta (5–7 Hz), alpha (8–12 Hz), beta (15–29 Hz), low gamma (30–59 Hz), and high gamma (60–90 Hz), using the open-source software, Brainstorm [110]. Each power band was then parcellated into 68 cortical regions [19].

### Dominance analysis

Dominance analysis seeks to determine the relative contribution (“dominance”) of each input variable to the overall fit (adjusted *R*^2^) of the multiple linear regression model (https://github.com/dominance-analysis/dominance-analysis [5, 17]). This is done by fitting the same regression model on every combination of input variables (2^*p*^*−* 1 submodels for a model with *p* input variables). Total dominance is defined as the average of the relative increase in *R*^2^ when adding a single input variable of interest to a submodel, across all 2^*p*^*−* 1 submodels. The sum of the dominance of all input variables is equal to the total adjusted *R*^2^ of the complete model, making total dominance an intuitive measure of contribution. Significant dominance was assessed using the spin test (see *Null models*), whereby dominance analysis was repeated between a spun disorder map and the original predictor matrix (1 000 repetitions).

Each multilinear model was cross-validated using a distance-dependent method proposed by [46]. Briefly, for each of 1 000 iterations, the 75% of regions closest in Euclidean distance to a randomly chosen source node were selected as the training set, and the remaining 25% of regions as the test set. Predicted values in the test set were then correlated to true abnormality patterns, and the correlations are shown in Fig. S3.

### Network spreading

Network spreading was computed as first introduced in Shafiei et al. [103] and later adopted in [23, 101]. Briefly, regional abnormality was defined as the normalized effect size used in all ENIGMA brain maps. For each region *i*, its neighbours are those with which region *i* is connected via a structural connection, as defined by the structural connectivity matrix. Mean neighbour abnormality of region *i* (*D*_*i*_) is the average abnormality of region *i*’s neighbours, where *d*_*j*_ represents the abnormality of neighbour *j*. Notably, this method normalizes neighbour abnormality by the number of connections made by region *i* (*N*_*i*_).

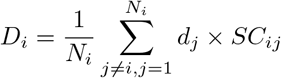

When neighbour abnormality is weighted by functional connectivity, each neighbour’s abnormality are weighted by the functional connection to node *i* (*FC*_*ij*_).

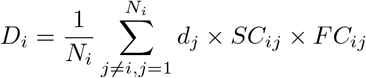

Each brain region was assigned a rank in terms of their node abnormality and their mean neighbour abnormality. The average of node and neighbour abnormality ranks was defined as the epicentre likelihood of the node, where nodes with high abnormality and whose neighbours are also highly atypical are more likely to be an epicentre of the disorder.

### Disorder similarity

For every brain region, we constructed a 13-element vector of disorder abnormality, where each element represents a disorder’s cortical abnormality at the region. For every pair of brain regions, we correlated the abnormality vectors to quantify how similarly two brain regions are affected across disorders. This results in a region-by-region matrix of “disorder similarity” (Fig. 5a). We verified that no single disorder pattern was driving the disorder similarity matrix by recalculating the disorder similarity when a single disorder is excluded. We then correlated the leave-one-out disorder similarity matrix with the original disorder similarity matrix. The minimum correlation was *r* = 0.95 (Fig. S9a). Finally, influence on the disorder similarity matrix by a disorder *i* was quantified as

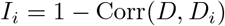

where *D* is the original disorder similarity matrix and *D*_*i*_ is the disorder similarity matrix constructed when disorder *i* is excluded (Fig. S9b).

### Null models

Spatial autocorrelation-preserving permutation tests were used to assess statistical significance of associations across brain regions, termed “spin tests” [2, 72]. We created a surface-based representation of the parcellation on the FreeSurfer fsaverage surface, via files from the Connectome Mapper toolkit (https://github.com/LTS5/cmp). We used the spherical projection of the fsaverage surface to define spatial coordinates for each parcel by selecting the coordinates of the vertex closest to the center of the mass of each parcel [121]. These parcel coordinates were then randomly rotated, and original parcels were reassigned the value of the closest rotated parcel (1 000 repetitions). Parcels for which the medial wall was closest were assigned the value of the next most proximal parcel instead. The procedure was performed at the parcel resolution rather than the vertex resolution to avoid upsampling the data, and to each hemisphere separately.

A second null model was used to test whether disorder similarity is greater in connected regions than unconnected regions. This model generates a null structural connectome that preserves the density, edge length, and degree distributions of the empirical structural connectome [11, 44, 94]. Briefly, edges were binned according to Euclidean distance. Within each bin, pairs of edges were selected at random and swapped. This procedure was then repeated 10 000 times. To compute a *p*-value, the mean disorder similarity of unconnected edges was subtracted from the mean disorder similarity of connected edges, and this difference was compared to a null distribution of differences computed on the rewired networks.

## Conflicts of Interest

CRKC, NJ, PMT received partial research support from Biogen, Inc., for research unrelated to this manuscript. JB has been in the past 3 years a consultant to/member of advisory board of/and/or speaker for Takeda/Shire, Roche, Medice, Angelini, Janssen, and Servier. He is not an employee of any of these companies, and not a stock shareholder of any of these companies. He has no other financial or material support, including expert testimony, patents, royalties. BF has received educational speaking fees from Medice GmbH. D.J.S. has received research grants and/or consultancy honoraria from Lundbeck and Sun.

## Acknowledgments

This research was undertaken thanks in part to funding from the Canada First Research Excellence Fund, awarded to McGill University for the Healthy Brains for Healthy Lives initiative. BM acknowledges support from the Natural Sciences and Engineering Research Council of Canada (NSERC Discovery Grant RGPIN #017-04265), the Canada Research Chairs Program, the Brain Canada Future Leaders Fund and the Healthy Brains for Healthy Lives initiative. JYH acknowledges support from the Helmholtz International BigBrain Analytics & Learning Laboratory, the Natural Sciences and Engineering Research Council of Canada, and the Fonds de reserches de Québec.

The research studies produced by the ENIGMA Working Groups would not be possible without the contributions of many researchers across the globe and the authors of this work thank all scientists who contribute to making this work possible. A full list of ENIGMA Consortium current and past members can be found here http://enigma.ini.usc.edu/ongoing/members/. The authors acknowledge the NIH Big Data to Knowledge (BD2K) award for foundational support and consortium development (U54 EB020403 to PMT) and support from NIMH R01MH116147 (PMT), NIMH R01MH116147 (JAT, TGMvE), NIMH R01 MH117601 (NJ, LS), NIMH R01MH085953 (CEB), NIMH R21MH116473 (CEB), NIMH 1U01MH119736 (CEB). For a complete list of ENIGMA-related grant support please see here: http://enigma.ini.usc.edu/about-2/funding. JB has been supported by the EU-AIMS (European Autism Interventions) and AIMS-2-TRIALS programmes which receive support from Innovative Medicines Initiative Joint Undertaking Grant No. 115300 and 777394, the resources of which are composed of financial contributions from the European Union’s FP7 and Horizon2020 Programmes, and from the European Federation of Pharmaceutical Industries and Associations (EFPIA) companies’ in-kind contributions, and AUTISM SPEAKS, Autistica and SFARI; and by the Horizon2020 supported programme CANDY Grant No. 847818). BF is supported by the European Community’s Horizon 2020 Programme (H2020/2014 – 2020) under grant agreements n° 667302 (CoCA), n° 728018 (Eat2beNICE), and n° 847879 (PRIME). This work was supported by a personal Veni grant to MH from the Netherlands Organization for Scientific Research (NWO, grant number 91619115). CRM is supported by NIH R01 NS065838; R21 NS107739. DJS is supported by South African Medical Research Council. GM is funded by a Wellcome Trust & The Royal Society Sir Henry Dale Fellowship [202397/Z/16/Z]. The funders had no role in study design, data collection and analysis, decision to publish or preparation of the manuscript.

---

**Figure S1.**
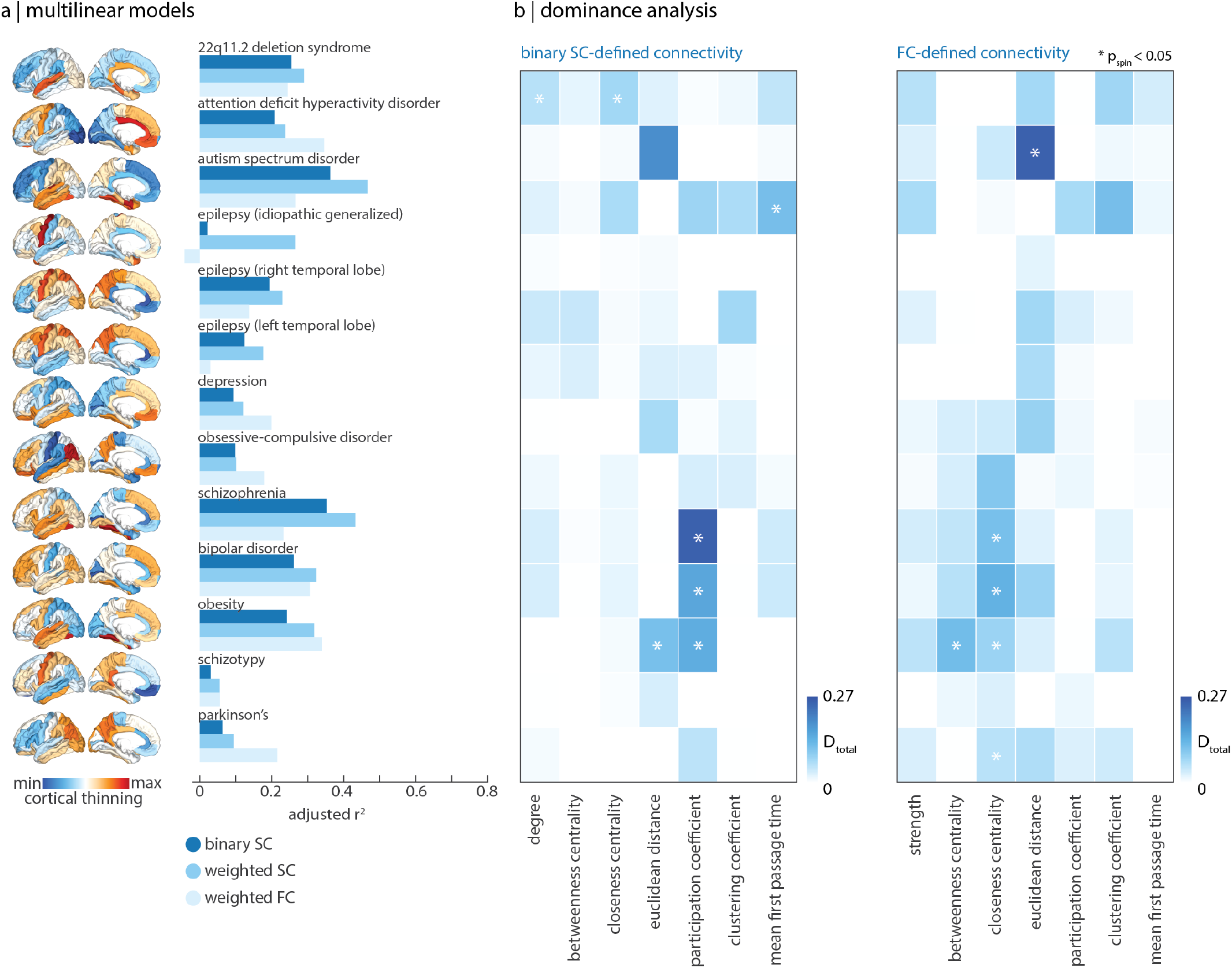
Computing global network measures on different connectivity matrices. Global network measures were calculated on the binary structural connectivity matrix as well as the weighted functional connectivity matrix. (a) Adjusted *R*^2^ between connectivity predictors and disorder maps (left-most surfaces) when connectivity measures were calculated on the binary structural connectome (dark blue), weighted structural connectome (medium blue; used in main text analyses), and weighted functional connectome (light blue). (b) Dominance analysis for the binary structural connectome (left) and weighted functional connectome (right). Asterisks represent *p*_spin_ < 0.05.

**Figure S2.**
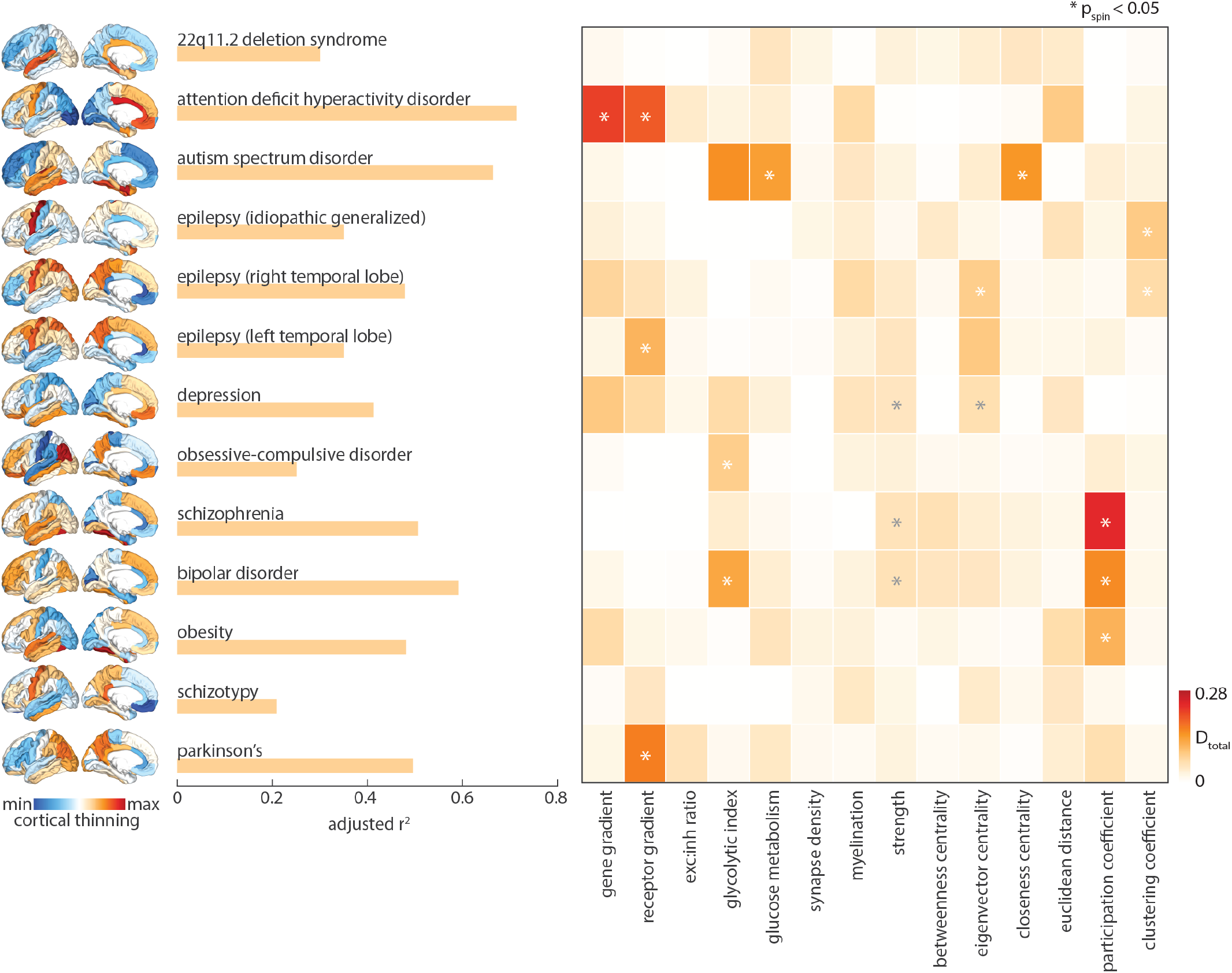
Combining molecular and connectomic predictors. Thirteen multivariate regression models were fit between the combined (molecular and connectomic) predictor set, to predict disorder-specific cortical thinning. Dominance analysis was applied to assess the predictors that contribute most to each mode. Asterisks represent *p*_spin_ < 0.05.

**Figure S3.**
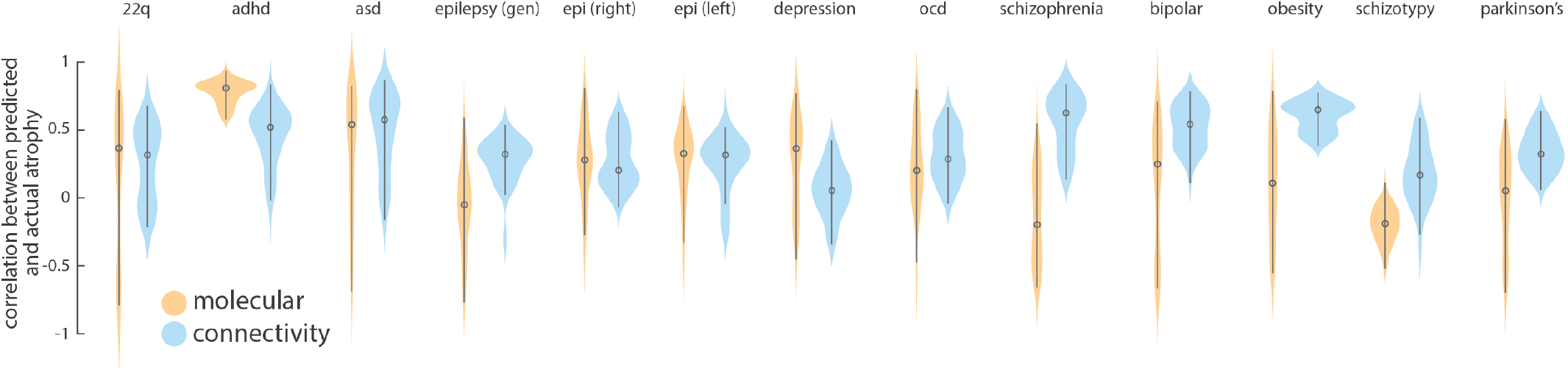
Distance-dependent cross-validation. Each model was cross-validated using a distance-dependent method. For 1 000 randomly chosen source regions, the model was fit on a training set, composed of the 75% of regions closes to the source region. Next, cortical abnormality values were predicted on the remaining 25% of regions and correlated to the empirical abnormality values. Circles represent the median and lines span the first to third quartiles.

**Figure S4.**
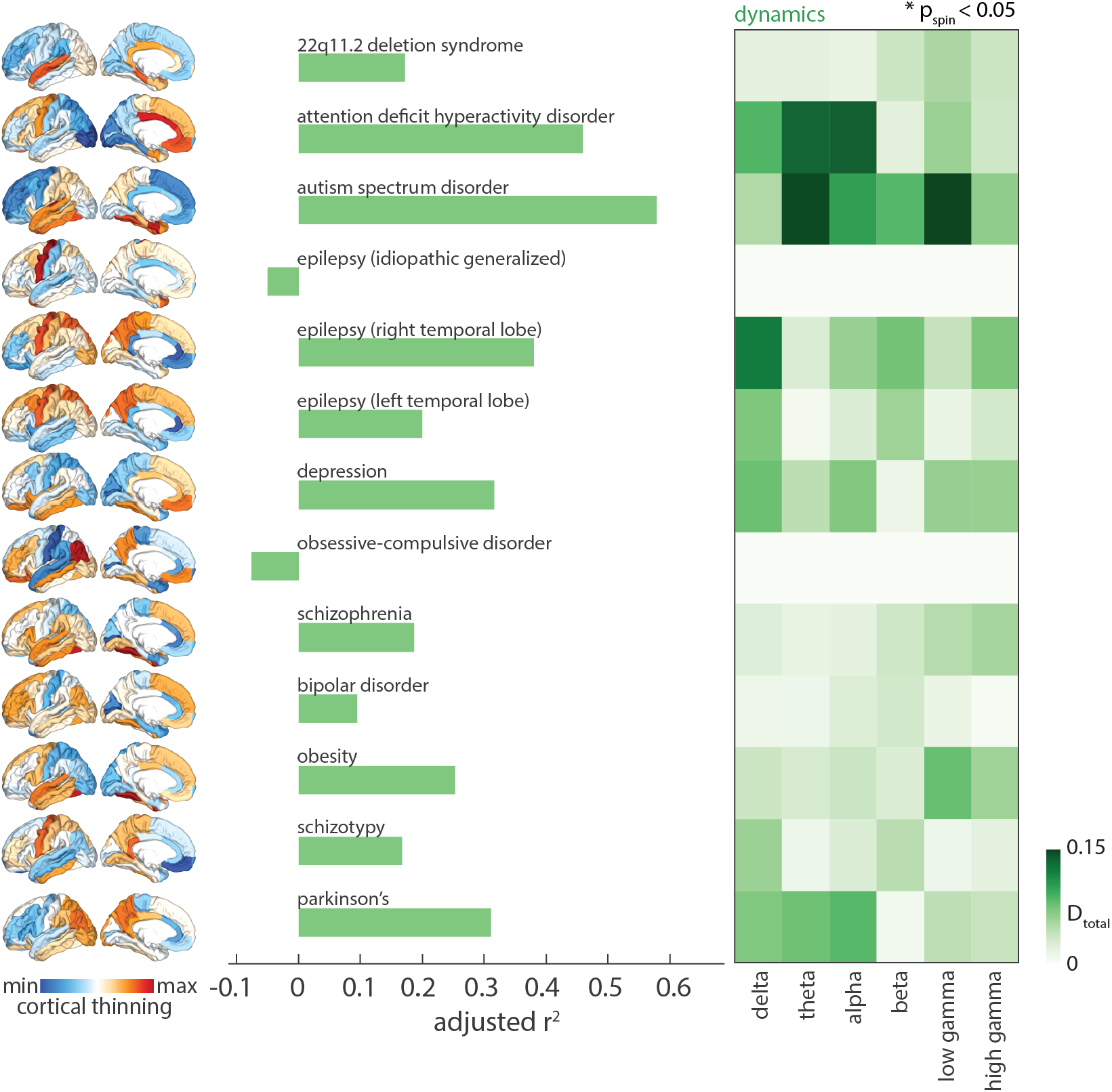
Mapping MEG-derived temporal predictors to disorder-specific cortical morphology. For each disorder, a multi-linear model was fit between six MEG-derived power distributions and the abnormality pattern. Dominance analysis extracted no significantly dominant predictors, and fits always underperformed compared to local biological predictors.

**Figure S5.**
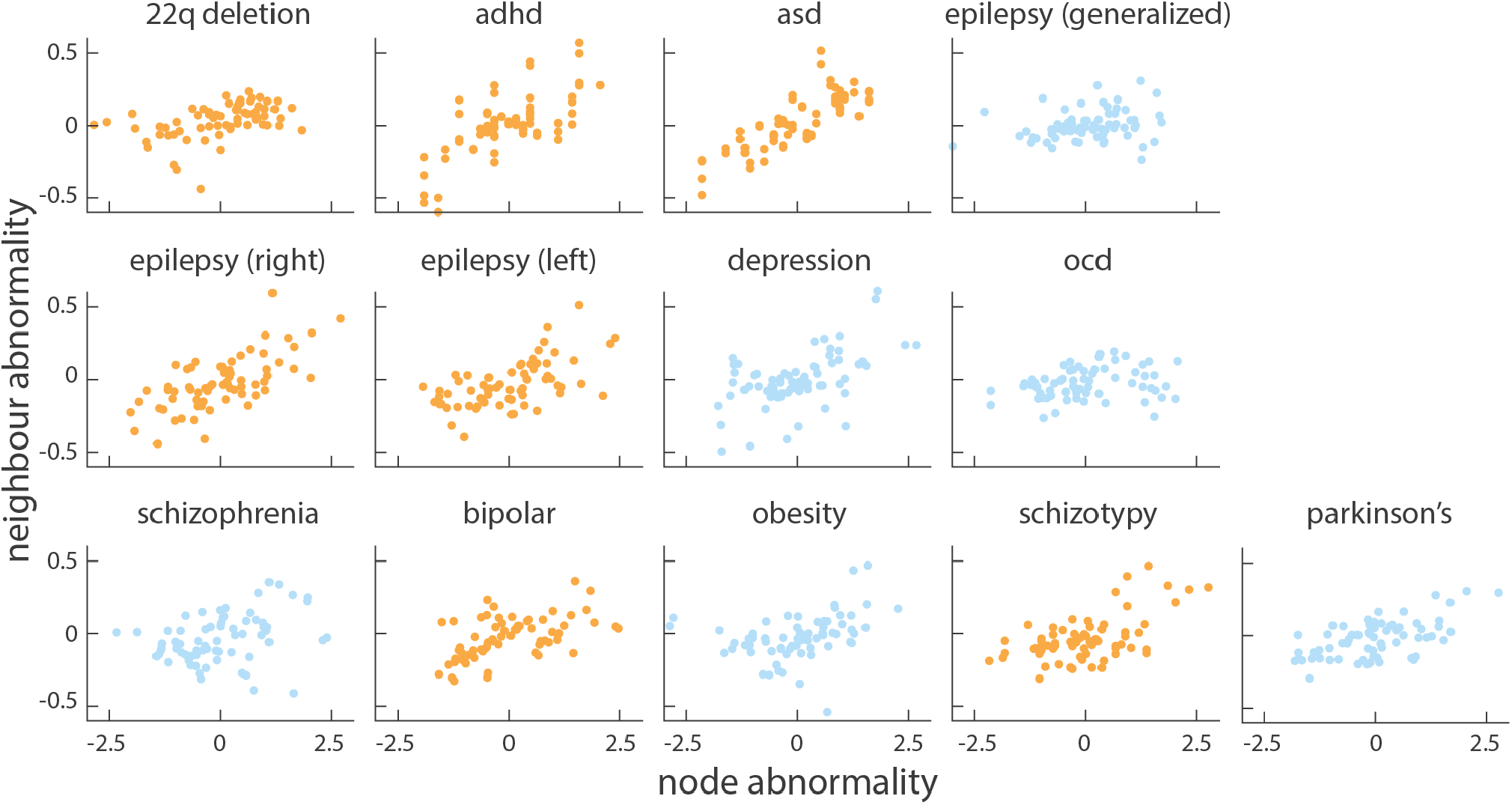
Assessing network spreading disorder-specific cortical morphology using structural and functional connectivity. A disorder whose cortical morphology demonstrates network spreading was defined as one whose regional abnormality pattern is correlated to mean neighbour abnormality, weighted by structural connectivity and functional connectivity. Yellow scatter plots indicate significant (*p*_spin_ < 0.05) node-neighbour correlations.

**Figure S6.**
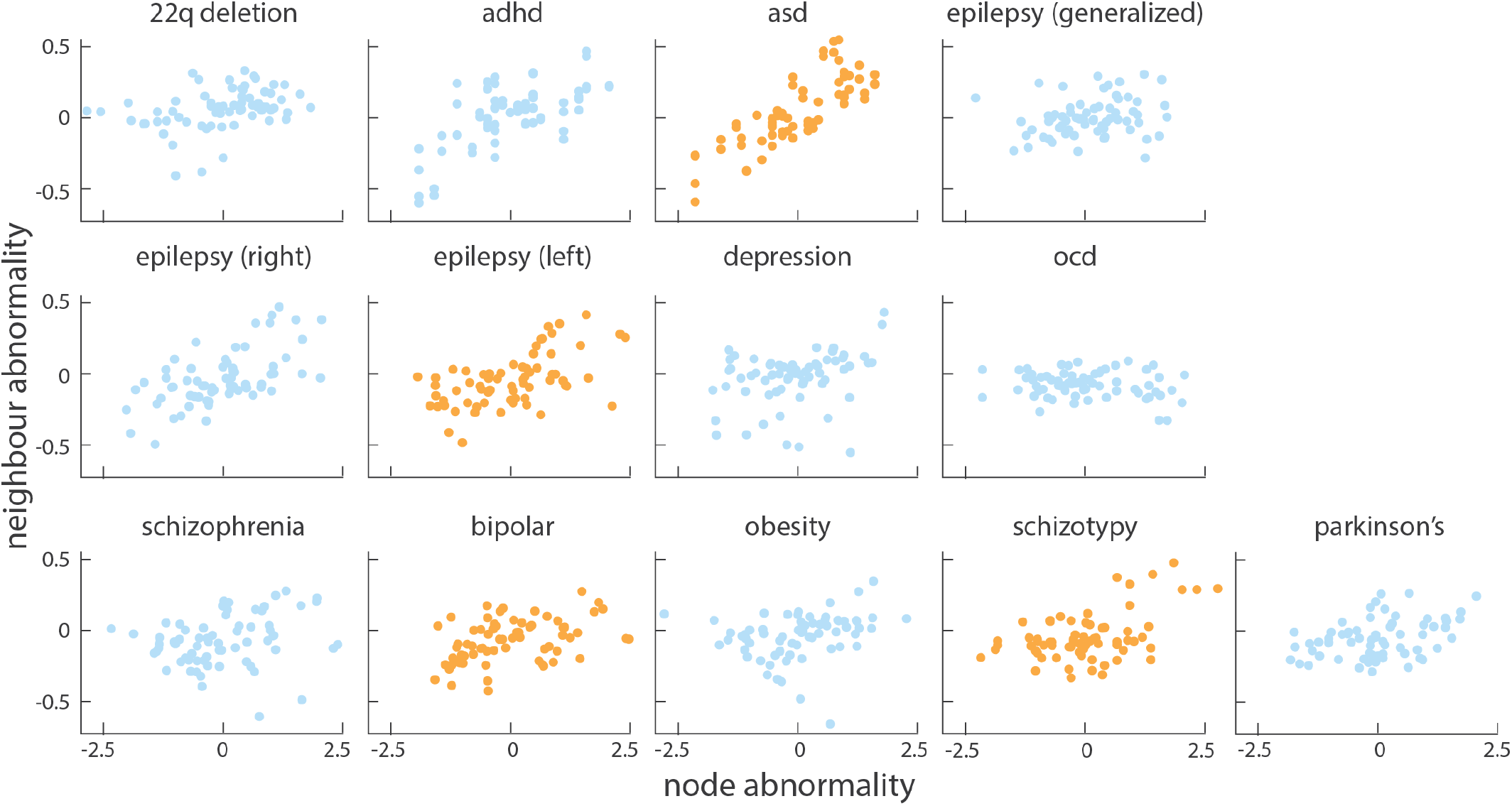
Assessing network spreading disorder-specific cortical morphology using structural connectivity only. A disorder whose cortical morphology demonstrates network spreading was defined as one whose regional abnormality pattern is correlated to mean neighbour abnormality, weighted by structural connectivity only. Yellow scatter plots indicate significant (*p*_spin_ < 0.05) node-neighbour correlations.

**Figure S7.**
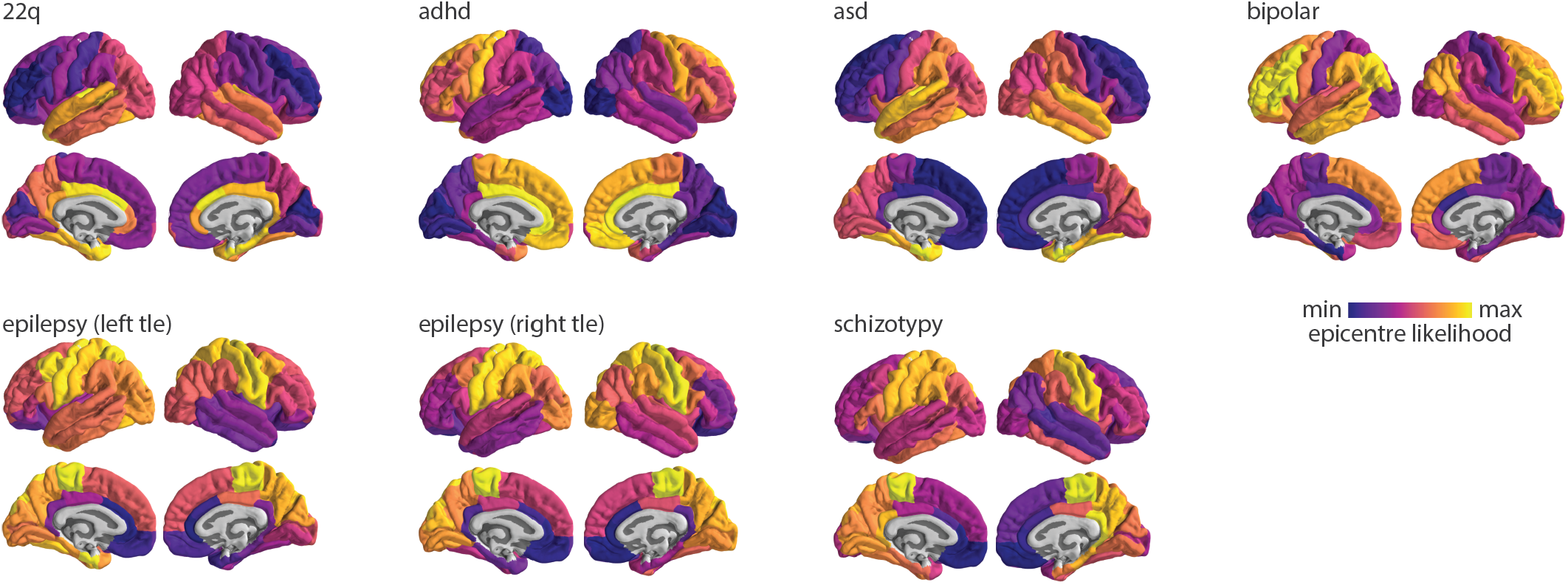
Epicentre likelihood. Epicentre likelihoods of the seven disorders that demonstrate significant correlations between node abnormality and mean sc- and fc-weighted neighbour abnormality.

**Figure S8.**
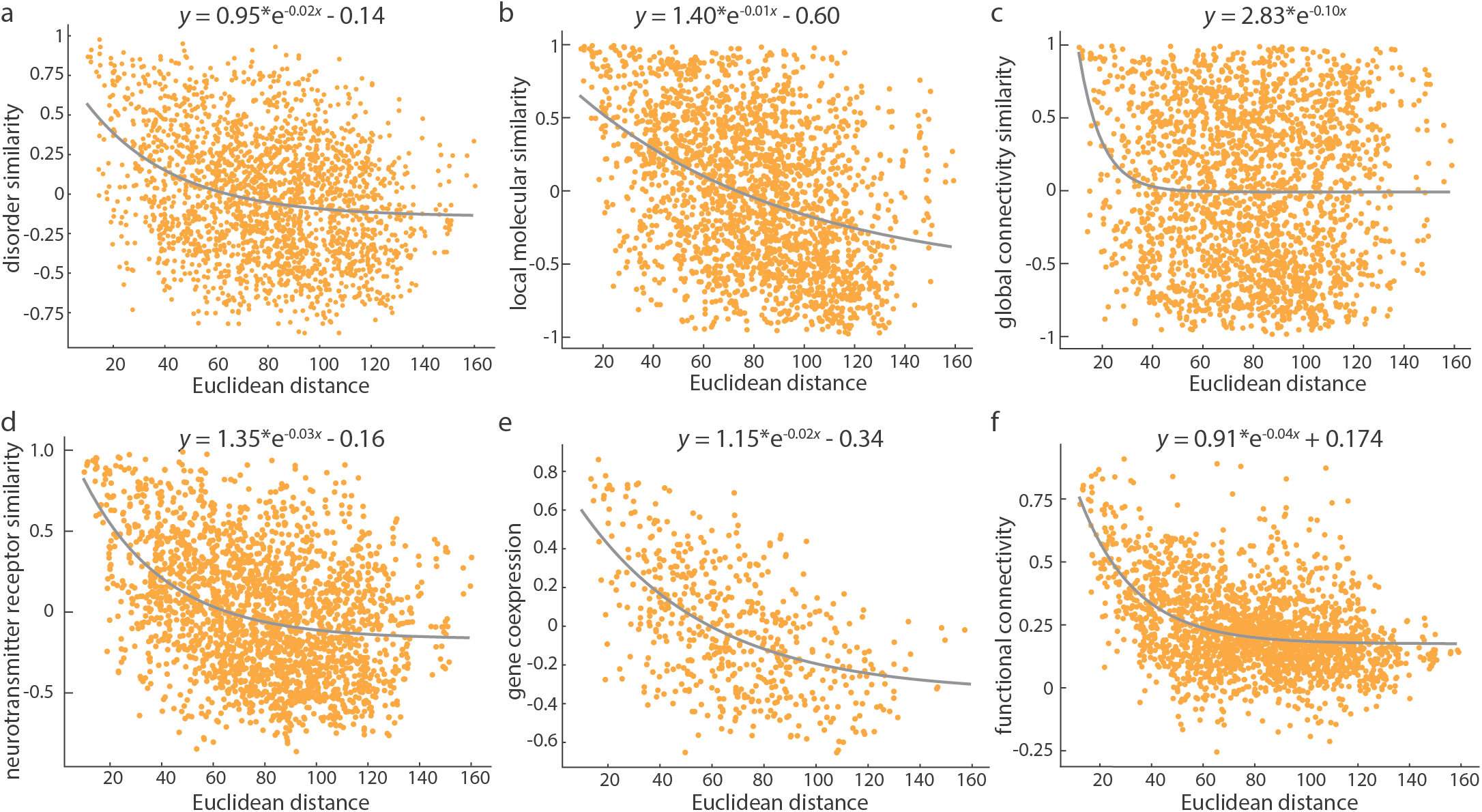
Annotation similarity between brain regions decreases exponentially. Similarity matrices for (a) disorder-specific cortical morphology, (b) local molecular fingerprint similarity, (c) global connectome predictor similarity, (d) neurotransmitter receptor density, (e) gene coexpression, and (f) functional connectivity were plotted against Euclidean distance. Each panel shows a negative exponential relationship between regional annotation similarity and distance, indicating that regions that are further apart show less similar properties. Note that gene coexpression was computed only for the left hemisphere.

**Figure S9.**
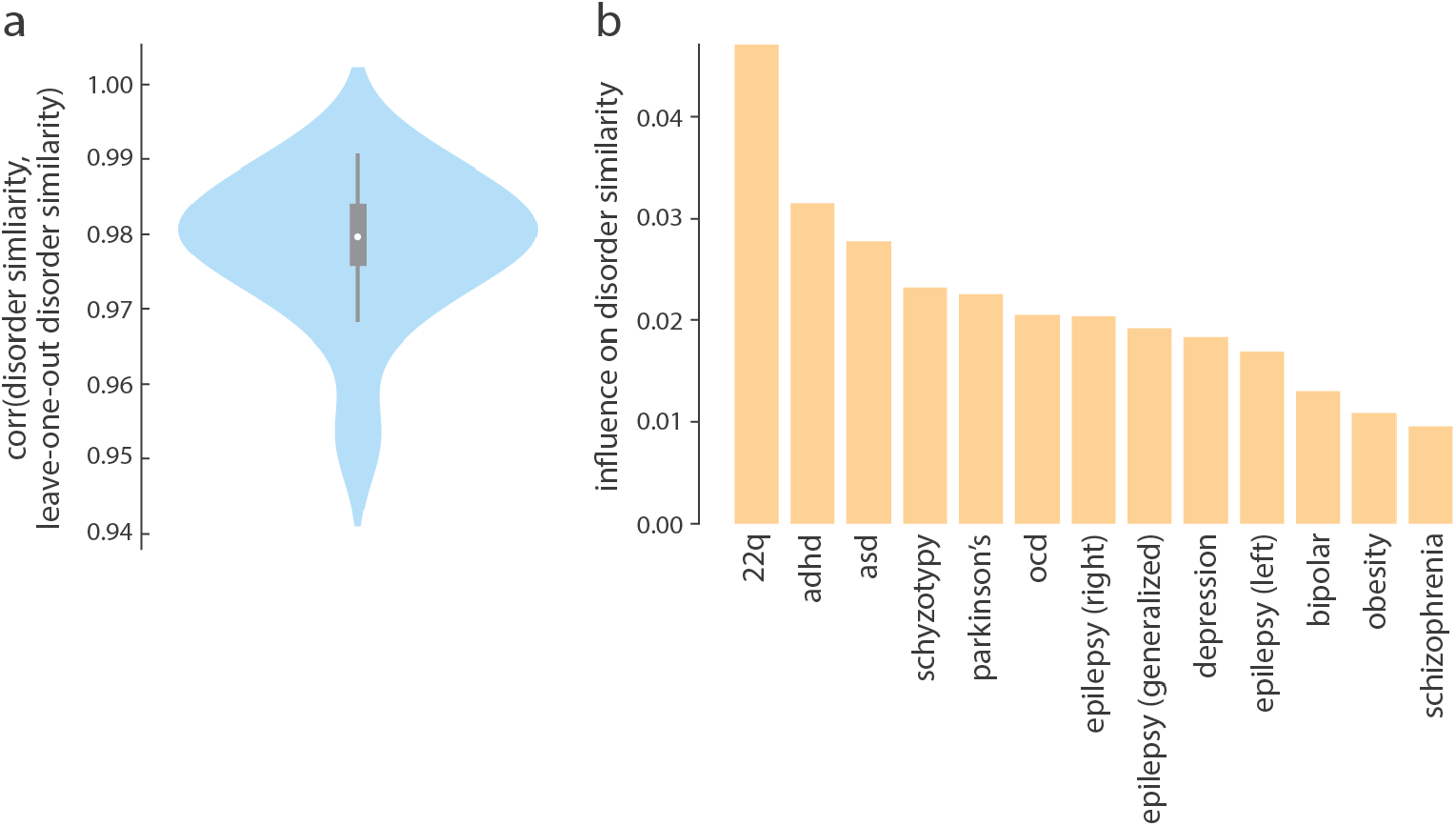
Robustness of disorder similarity matrix. (a) Disorder similarity was correlated to a version of the disorder similarity matrix, constructed by excluding a single disorder, across all thirteen disorders. The minimum correlation, computed when 22q-deletion syndrome is excluded, is *r* = 0.95. (b) The influence that each disorder has on the disorder similarity matrix is calculated as the difference between 1 and the correlation coefficient calculated in (a).

## Notes

https://github.com/netneurolab/hansen_crossdisorder_vulnerability

### Competing Interest Statement

The authors have declared no competing interest.

